# Imaging intracellular components *in situ* using super-resolution cryo-correlative light and electron microscopy

**DOI:** 10.1101/2023.11.19.567713

**Authors:** Mart G. F. Last, Lenard M. Voortman, Thomas H. Sharp

**Affiliations:** Department of Cell and Chemical Biology, Leiden University Medical Centre, 2300 RC Leiden, The Netherlands

**Keywords:** Super-resolution light microscopy, cryoEM, cryo-electron tomography, correlative light-electron microscopy, CLEM, SR-cryoCLEM, superCLEM

## Abstract

Super-resolution cryo-correlative light and electron microscopy (SRcryoCLEM) is emerging as a powerful method to enable targeted *in situ* structural studies of biological samples. By combining the high specificity and localization accuracy of single-molecule localization microscopy (cryoSMLM) with the high resolution of cryo-electron tomography (cryoET), this method enables accurately targeted data acquisition and the observation and identification of biomolecules within their natural cellular context. Despite its potential, the adaptation of SRcryoCLEM has been hindered by the need for specialized equipment and expertise. In this chapter, we outline a workflow for cryoSMLM and cryoET-based SRcryoCLEM, and we demonstrate that, given the right tools, it is possible to incorporate cryoSMLM into an established cryoET workflow. Using Vimentin as an exemplary target of interest, we exemplify all stages of an SRcryoCLEM experiment: performing cryoSMLM, targeting cryoET acquisition based single-molecule localization maps, and correlation of cryoSMLM and cryoET datasets using scNodes, a software package dedicated to SRcryoCLEM. By showing how SRcryoCLEM enables the imaging of specific intracellular components in situ, we hope to facilitate the further adaptation of the technique within the field of cryoEM.

## Introduction

Cryogenic correlated light and electron microscopy (cryoCLEM) has enormous potential as a method for *in situ* imaging studies on biological samples^1,2^. By combining the specificity of fluorescent labelling with the high resolution of cryo-electron microscopy (cryoEM), cryoCLEM enables various advanced applications, such as targeted cryoEM data acquisition or targeting lamella preparation by focused ion beam^3^; applications that have hitherto been very challenging by use of electron microscopy alone.

A promising approach to cryoCLEM is to combine super-resolution fluorescence microscopy with cryo-electron microscopy, usually cryo-electron tomography (cryoET). The aim of using super-resolution rather than widefield fluorescence microscopy is to attain a fluorescence imaging resolution that is sufficiently high to enable not just targeted cryoEM acquisition on single biomolecules or complexes, but also the identification of unknown biomolecules that are visible within cryoET datasets on the basis of accurately correlated single-molecule annotations^4^.

Widespread uptake of super-resolution cryoCLEM (SRcryoCLEM) has been limited, in part, due to the need for dedicated hardware, as well as the steep learning curve that is often associated with novel methods. Since, in our experience, SRcryoCLEM can be routinely and straightforwardly applied to study various biological systems, we seek to promote adaptation of the technique by reporting advancements in hardware, software, and methodology.

In this chapter, we describe a full workflow for super-resolution cryoCLEM (SRcryoCLEM), including; cell culture, cryosample preparation, cryoSMLM acquisition and processing, cryoET acquisition and tomogram reconstruction, correlation, fluorescence guided particle picking, and finally protein structure determination by sub-tomogram averaging. We demonstrate each of these steps using Vimentin as the target protein, and rsEGFP2-Vimentin expressing U2OS cells cultured on EM grids as the sample.

Many of the steps employed in our workflow have already been described in detail in various other publications^5,6^. The focus of this chapter is therefore on the challenges and opportunities that are unique to SRcryoCLEM, such as cryoSMLM acquisition, targeted cryoET acquisition, and correlation of cryoSMLM and cryoET data. Where possible, we refer to earlier publications and protocols for more standard procedures but also provide more detailed descriptions of specific steps in the procedure that could not be included in those protocols.

Practically, combining cryo-SMLM and cryoET is mostly a matter of incorporating cryoSMLM into a previously established cryoEM workflow. In our lab, we do so using the ‘cryoscope’, a custom fluorescence microscope described in another chapter of this book, but commercial fluorescence cryo-microscopes are also available. Examples include various upright light microscopy systems that can be equipped with a Linkam CMS196 cryostage and the dedicated cryo-microscopes with a proprietary cryostage by Leica. While these are not necessarily suited for SMLM-type imaging, the data produced by such systems can alternatively also be used in the workflow described here. Many of the other methods outlined in this chapter, e.g., those for correlation, can also be applied when using these microscopes, but in this chapter we focus on the use of the cryoscope. Where relevant, alternatives to methods specific to the cryoscope are suggested.

## Methods

### Materials and equipment

The following pieces of equipment are required for this workflow. Standard consumables that are a part of their use, e.g., tweezers, pipette tips and gloves, are not listed.

- Glow discharger (here: PELCO easiGLOW)
- Cell culture hood & incubator suited for working with mammalian cells
- Plunge freezer - one that can be used to blot grids from the back only (here: Leica EM GP)
- Hot plate
- Cryoscope^1^ or any equivalent cryo-microscope.
- Windows or Linux PC with a CUDA enabled GPU, to run scNodes.
- CryoEM suited for tilt series acquisition (here: Talos Arctica, Thermo Fisher Scientific)

The following materials are used in the workflow.

- 3.5 mm cell culture dishes (here: Corning #430165)
- 1.2/1.3 300 mesh UltrAuFoil holey gold TEM grids (QuantiFoil)
- Sterile phosphate buffered saline solution
- Cell culture medium (here: Dulbecco’s Modified Eagle Medium (DMEM), supplemented with 10% fetal bovine serum and 100 units/mL of penicillin-streptomycin.
- The cell line of interest
- Autogrid rings and C-Clips, depending on the model of cryoEM used

### Sample preparation

CryoEM requires thin samples, and there are various ways of obtaining sufficiently thin samples, the choice of which to use depends on what structure one desires to image^7^. For certain cytosolic components, ubiquitous organelles such as endosomes and lysosomes, or the cell membrane - i.e., structures that can be found in the periphery of cells – adherent cells cultured on EM grids provide ample suitably thin areas to enable direct imaging. For structures that are rare or not found in naturally thin regions of cells, thinning by focused ion beam (FIB) milling is one commonly applied strategy^8^, but alternative methods such as cryo-ultramicrotomy, microsome formation^9^, or gentle purification of organelles have also found use^10,11^.

The requirements that cryoSMLM imposes on the sample are of a different nature, and generally do not limit experimental throughput as much as the requirement of sample thickness does. CryoSMLM relies on the labelling of the structure of interest with a fluorescent probe that can be photo-modulated, i.e., turned on or off in a controlled manner by exposure to different light sources. A growing number of fluorescent proteins that can be used for cryoSMLM are known^12-15^; here, we use the green reversibly photo-switchable fluorescent protein rsEGFP2^16^.

The type of EM grid that is used is another important parameter in cryoSMLM, and dictates the maximum fluorescence excitation light intensity that can be used. We and others have previously reported on this topic, and we refer to those reports for more in-depth information^17,18^. Here, we use 1.2/1.3 UltrAuFoil 300 mesh holey gold grids, which are commercially available and well suited for cryoSMLM in general, and especially for adherent mammalian cells.

To target Vimentin, we made use of a genome-edited U2OS cell line that expressed endogenously labelled Vimentin-rsEGFP2, that was originally prepared by Ratz et al.^19^ Endogenous labelling has the benefit of conserving the endogenous expression levels of the protein, leading to a more homogenously labelled sample with a generally lower label density and fewer expression-induced artefacts in comparison to cells that are stained *via* overexpression.^19^ A useful and detailed protocol on culturing cells on EM grids, including information on inducing expression of fluorescently labelled proteins, is that by Hampton et al.^20^

#### Glow discharging grids

- Carefully move four (or more) grids into a 3.5 cm cell culture dish, with the support film facing up. Although the dish will be sterilized later, it is best to wear gloves to avoid contaminating the dish.
- Mount the culture dish (without the lid) in the glow discharger and glow discharge for 60 seconds at 25 mW.

#### Sterilizing grids

- Move the dish into a cell culture hood.
- Carefully deposit 1 mL of sterile phosphate buffered saline (PBS) in the dish. The grids should be submerged. If a grid floats, carefully pick it up and force it under.
- Turn on the UV sterilization lamp and leave the dish, without the lid, to sterilize for at least 30 minutes.

#### Cell culture

- Carefully remove the PBS from the culture dish using a sterile vacuum aspirator.
- Add 2 mL of pre-heated (37 °C) culture medium (here: DMEM supplemented with 10% fetal bovine serum and 100 units/mL of both penicillin and streptomycin) to the dish. If a grid floats, carefully force it under again.
- Seed ∼100,000 cells in the dish, so that the expected number of cells per grid square is around one.
- Leave the dish to incubate in a culture cabinet overnight, at 37 °C and in 5% CO_2_.

#### Plunge freezing

- Turn on the hot plate and set the target temperature to 37 °C. Place 10 mL of PBS on the hot plate in a small vial or dish.
- Turn on the plunge freezer and set the humidity to 99% and temperature to 37 °C. Wait for ∼30 minutes for the chamber to reach these conditions.
- Cool down and initialize the plunge freezer. Set the blotting time to 4 seconds. Place a cryo grid box in the designated slot, for storage of grids after vitrification.
- Move the culture dish from the culture cabinet onto the hot plate, which should now be at temperature.

The following procedure is repeated for every grid.

- Pick up one grid out of the culture dish with a plunge tweezer and gently dip it into the vial of PBS to wash away the culture medium. Repeat two more times; the culture medium is fluorescent, and so it helps to remove it. Be careful only to submerge the grid and the very tip of the tweezer, in order to not wet the tweezer too much.
- Mount the tweezer in the plunge freezer, such that the cells face away from the blotting paper (if possible).
- Blot and plunge the grid and store it in the grid box.

Depending on the model of cryoEM that is used, grids may need to be mounted into a c-clip (‘clipped’) after plunge freezing. We prefer to do this right after plunging in order to avoid spending time imaging sites that may otherwise subsequently be damaged during clipping. It is practical to choose a standard orientation of the grid during clipping, with the cells either facing down or up, in order to be aware in which orientation samples are mounted in the light microscope.

### CryoSMLM

Accurate localization of single molecules is best done with bright fluorophores against a dark background. Unfortunately, a number of complications arise in low temperature conditions. One is that numerous endogenous chemical compounds, such as NADH and FAD, emit notable fluorescence in cryogenic conditions^21^, and thus contribute to the background signal. A second complication is that the photo-conversion kinetics of fluorescent proteins are significantly reduced at a low temperature^12^, and that as a result the extent of de-activation of the population of fluorophores is limited. Third, the efficiency of photo-activation is also limited^12^. Thus, a high illumination intensity is required in order to activate a sufficient number of fluorophores, but at the same time the intensity must also be limited in order to avoid heating the sample.

These effects can be particularly problematic in densely labelled or thick areas of the cell, such as near the nucleus, where the background fluorescence intensity is often brighter than the single molecule events. Fortunately, imaging in thin regions of the sample is a convenient way of avoiding this problem: peripheral regions of the cell with a thickness optimal for cryoEM are also optimal for cryoSMLM, as the depth of focus of a microscope exceeds the sample thickness in these areas.

#### Sample loading

The instructions below are written with the use of a Linkam Scientific ‘CMS196v3’ cryostage in mind, but are similar for other cryostages.

- Make sure that the cryostage sample mount is open and in the loading position. Power the stage, then launch the microscope control software.
- Cool down the cryostage by filling the dewar with liquid nitrogen. Turn on the hot plate and set it to 45 °C. After usage, always leave cold tools on the hot plate to heat and dry.
- Once the LN2 dewar is cold and full, press the button to start cooling the sample chamber.
- Wait ∼15 minutes for the chamber to cool down fully. Use the LN2 dewar and lid stops to prevent moisture accumulating within the cold chambers. These stops should be in place whenever the stage is not actively used or in the microscope, in order to minimize contamination.
- Using cooled tweezers, carefully but quickly transfer the grid box that contains the samples into the cryostage chamber.
- Use the grid box opening tool to nudge the grid box into its designated slot, then open the grid box (Fig. 1a).
- Using fresh tweezers, mount a grid in the cryostage sample mount (Fig. 1b). At this point, make sure that the grid is oriented in the desired manner (see below). Close the mount and the grid box.
- Place the mount on the bridge of the cryostage (Fig. 1cd), then turn the lid so that the opening is facing away from the grid.
- Mount the cryostage in the stage mounting clamp and move the vertical translation stage up, until the objective lens almost touches the stage lid. At this point, rotate the lid and carefully move the stage further upwards into the final position, which is when the sample is approximately 4 mm under the focal plane of the objective lens. Tighten the clamps on the translation stage.
- Turn on the stage’s condenser LED (via either the microscope control software or using the controls on the stage itself). Using the software, slowly move the objective lens down towards the sample while acquiring live images (the intensity should be seen to increase), until the sample appears in focus. If after ∼5 mm of movement no sample is visible, do not move the lens down any further – instead, gently lift the lid of the cryostage to inspect whether the sample is mispositioned or missing. *The bridge of the cryostage is very fragile and will be damaged by overextension of the lens*.

**Figure 1.**
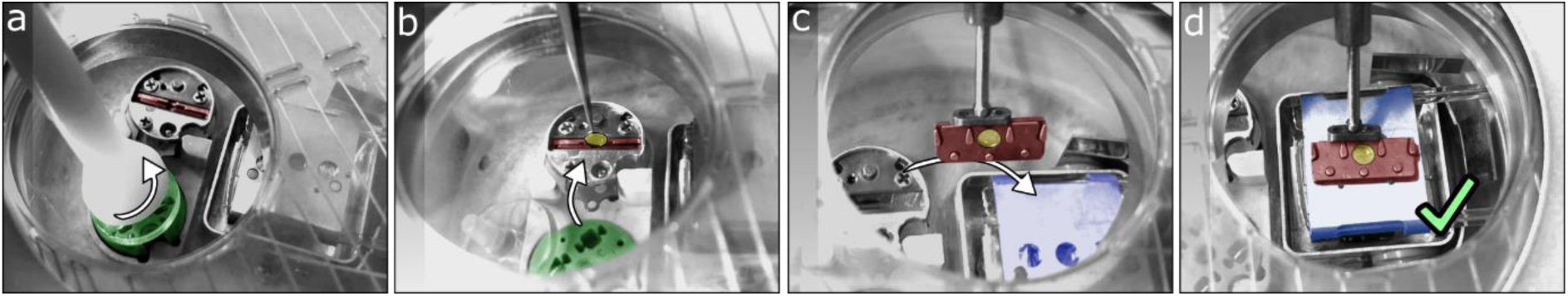
Handling grids in the Linkam CMS196 v3 cryostage. **a)** After cooling down the chamber cryo gridbox (green) is placed in the chamber and opened. **b)** Using cooled tweezers, a grid (yellow) is carefully picked up from the gridbox and placed in the sample holder (red). **c)** The sample holder is then transferred onto the bridge (blue) and **d)** placed such that the magnets in the holder snap onto the corresponding magnets on the bridge. The rotating lid of the stage can then be turned so that sample is not directly underneath the opening and be closed off with a stopper. After sealing the chamber against water contamintion *via* convection in this manner, the stage can be moved.

#### Selecting suitable sites for imaging

While it is generally recommended to load a sample into a microscope with the top facing up, the choice is yours in cryoSMLM. When the sample is loaded with the cells facing down, i.e., away from the objective lens (Fig. 2a), the material of the EM grid partially blocks light from exciting fluorescence in those regions of the sample that are directly on top of the grid bars or the solid area of the support film and that would thus not be suited for cryoEM data acquisition. This can be a convenient way to pre-screen locations. Moreover, the bottom side of cells grown on an EM grid is flatter than the top side, which can sometimes be highly curved and cause lensing artefacts. The reduced stray fluorescence and increased flatness can therefore positively affect the image quality.

**Figure 2.**
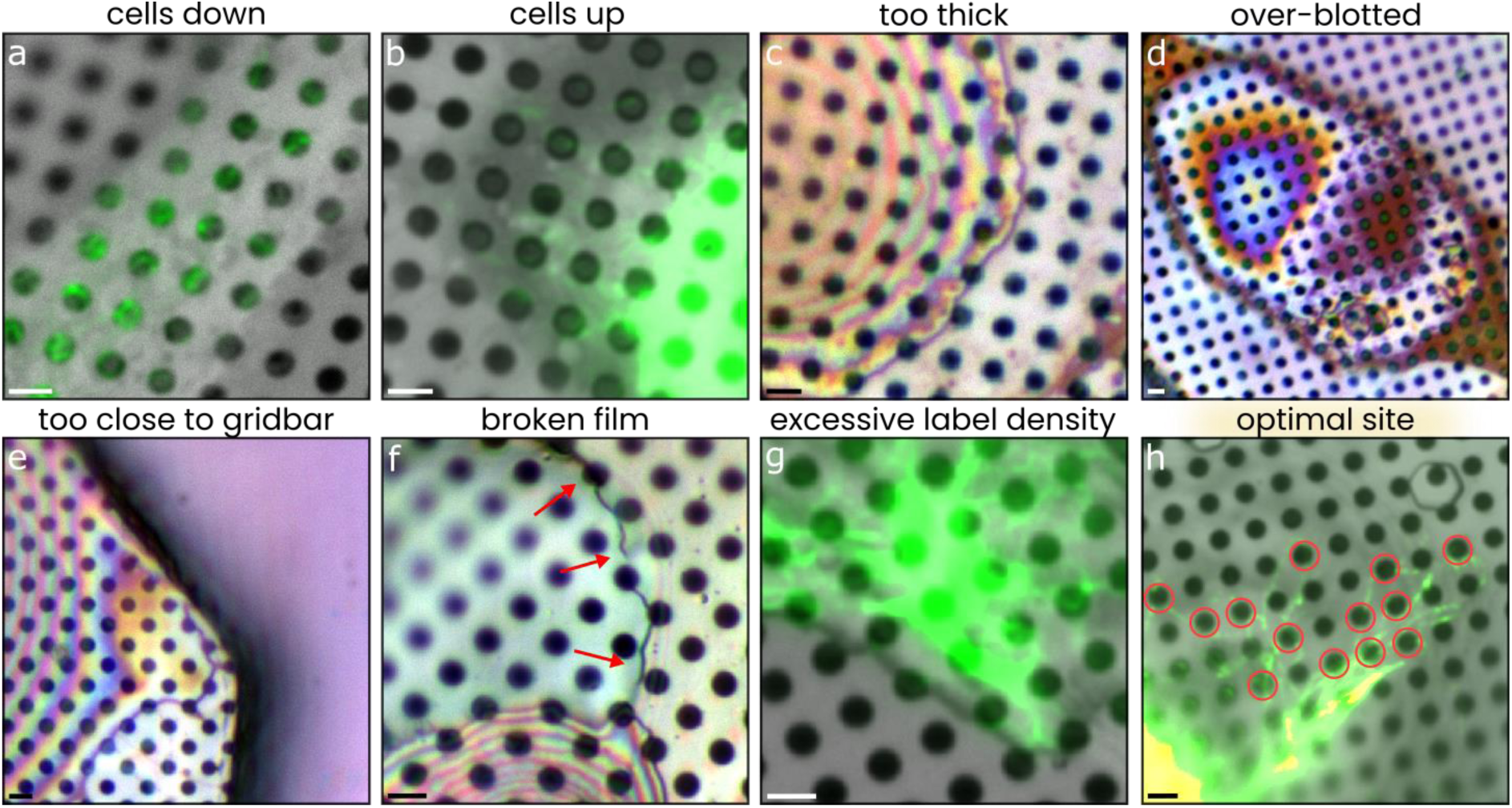
examples of sites that have useful or unsuitable characteristics for cryoSMLM. Panels show either reflected light (gray) and fluorescence (green), or composite reflected light (rgb) images that can be used to assess the sample thickness. In these latter examples, we have not applied the neural network described in reference ^22^, but rather gauge the thickness by eye, based on the composite reflection colour. **a)** A grid mounted upside down, so that the (gold) support film blocks excitation of fluorescence in regions directly on support film. **b)** A sample mounted with the cells facing up. The whole cell is now visible, enhancing context, but bright fluorescence from regions not suited for EM is also imaged. **c)** A cell with a relatively compact morphology, too thick for EM imaging. **d)** A cell that appears to be exceedingly thin due to excessive blotting (30 seconds in this example). When imaged by EM, these sites appear thoroughly destroyed. **e)** A suitably thin and flat region of a cell, but that is situated close to a grid bar and is thus not suited for tomography. **f)** Samples on damaged film are more prone to devitrification by illumination, as discontinuities in the film limit cooling, and are susceptible to beam induced drift in EM. **g)** High label densities lead to a bright background and to the presence of (many) simultaneously active, overlapping fluorophores that cannot not be individually resolved. **h)** A site optimally suited for both cryo SMLM and cryo EM: the region is thin, the labelling density moderate yet sufficient to initially resolve structures in single, wide-field images, and the structures of interest are also found within holes, on an intact film, away from the grid bars. The thirteen red circles indicate holes that would be of interest for high - resolution imaging. Scale bars are 2 µm.

When samples are loaded with the cells facing up (Fig. 2b), the fluorescent signal coming from within holes in the support film is far dimmer than that coming from adjacent regions that are on top of the solid area of the support film. However, the sample can now be observed in its entirety, including parts of cells that have grown on grid bars that could otherwise not be seen. In our experience, the context of the rest of the cell surrounding a small region of interest is an important indicator of whether the cell was intact and healthy. Since thin regions of the sample also tend to be sufficiently flat, we usually load samples facing up.

The quality of the sample and of the sites selected for imaging is the most important factor that influences the success of an SRcryoCLEM experiment. Provided the sample itself is well prepared, the first and most critical step of the cryoSMLM part of the workflow is thus to select regions of interest.

The single most important parameter dictating whether a site is worth imaging is the local sample thickness. This thickness can be estimated in a number of ways, including fluorescence, reflected light, and transmitted light imaging, optionally in combination with optical sectioning. On our microscope, we use a reflected light imaging method with which we can effectively generate an image of the sample thickness^22^. This rapid screening method allows us to filter out any sites that would not be suited for cryoEM (Fig. 1c).

A second, key parameter is the integrity of the cell and its positioning within the sample. When using a plunge freezer to vitrify samples, variability in the effect of blotting can lead to some cells being excessively blotted (Fig. 2d). These cells can appear ideally thin and often do not exhibit markedly different patterns of fluorescence, but upon observation in the cryoEM present themselves as cellular debris devoid of recognizable structure. Regions of cells that have grown close to a grid bar (Fig. 2e) are also impractical, as the grid bars limit the tilt range in cryoET, as are sites on or close to tears in the support film (Fig. 2f), as discontinuities in the film limit the dissipation of heat (generated by absorption of light), thus leading to devitrification, as well as cause increased drift in cryoEM.

Finally, sites with a high label density are also to be avoided, as the low efficiency of photo-deactivation of RSFPs leads to high background fluorescence or to the simultaneous presence of multiple, overlapping fluorescent particles that are not individually resolvable (Fig 2g).

#### Screening the sample

With that in mind, the next step is to locate sites of interest that are suited for both cryoSMLM and cryoET (Fig. 3).

- In the microscope software, set up a live feed with two imaging channels: 1) reflected light, and 2) rsEGFP2 fluorescence.
- Browse the sample and look for suitable sites, keeping in mind the points of attention illustrated in Figure 1.
- When a site is found, save its position, and then continue searching; the first site is rarely the best.
- After locating a sufficient number of suitable sites, find the center marker of the grid and note down this stage position as well. This can be of help when relocating sites in the electron microscope.

**Figure 3.**
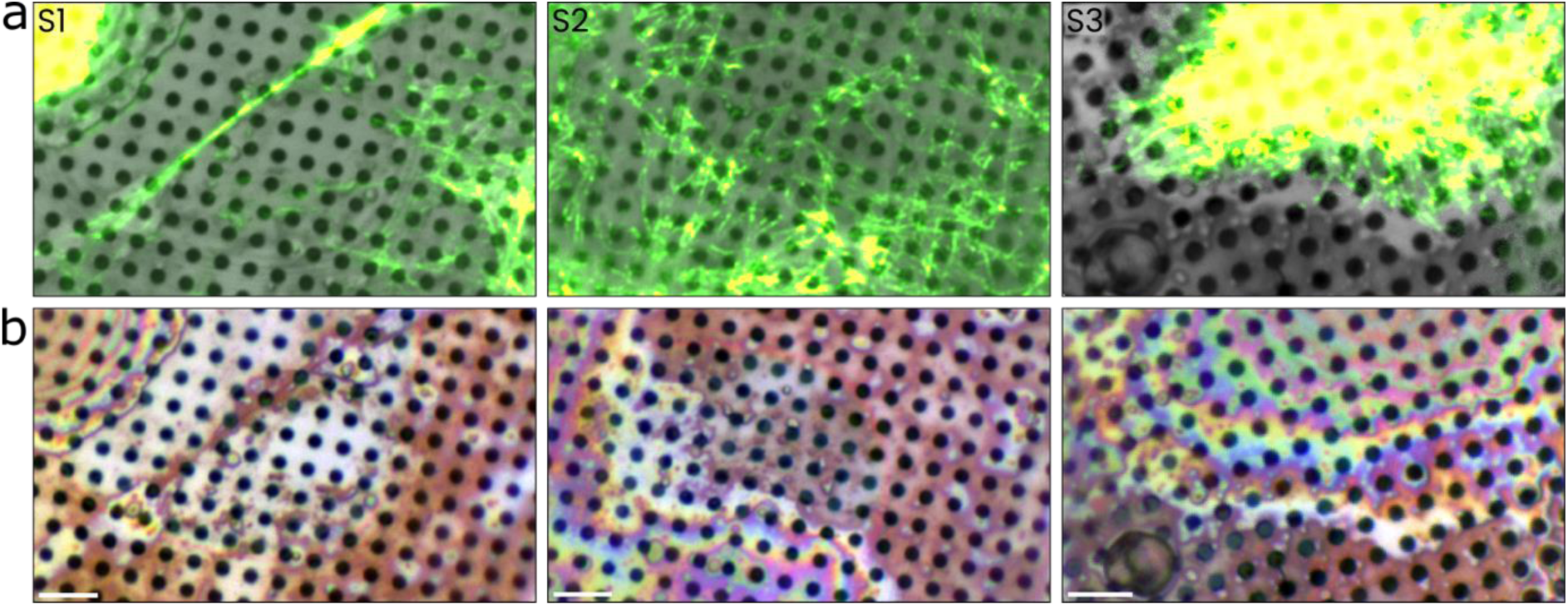
Sites of interest selected for SR-cryo CLEM imaging. **a)** Widefield images of rsEGFP2 - Vimentin fluorescence (green) overlayed on reflected light images (gray), in the sites (S1, S2, S3) selected for further imaging. **b)** Composite reflected light images of the same sites, indicating that the thickness of these regions is sufficiently low to enable high - resolution cryo ET data acquisition. Scale bars are 5 µm.

#### SMLM acquisition

After locating suitable regions of the sample, the next step is to acquire fluorescence timelapses that can be used for single-molecule localization. As stated before, this requires the diffraction limited signal from individual fluorescent molecules to be clearly identifiable over background fluorescence and noise.

In thermal equilibrium, a population of rsEGFP2 fluorophores is partially in both the fluorescent on-state as well as in the dark off-state. When the local label density is high, the overall fluorescent signal is consequently bright, impeding the ability to resolve individual molecules.

To take this issue in to account, we generally use two slightly different illumination routines: we begin, if necessary, with a ‘depletion’ routine, followed by a ‘switching’ acquisition. A depletion acquisition (Fig. 4, Table 1) is intended to reduce the overall brightness and simply involves prolonged fluorescence excitation using 488 nm light, a wavelength that excites fluorescence as well as deactivates rsEGFP2. Typically, we use a total depletion time of 100 seconds (1000 frames x 100 ms exposure, ∼180 W/cm^2^), after which the fluorescence intensity is reduced to less than 20% of the initial value. Towards the end of such an acquisition, single-molecule blinking can usually be observed (due to a low rate of photoactivation of rsEGFP2 by 488 nm light).

**Figure 4.**
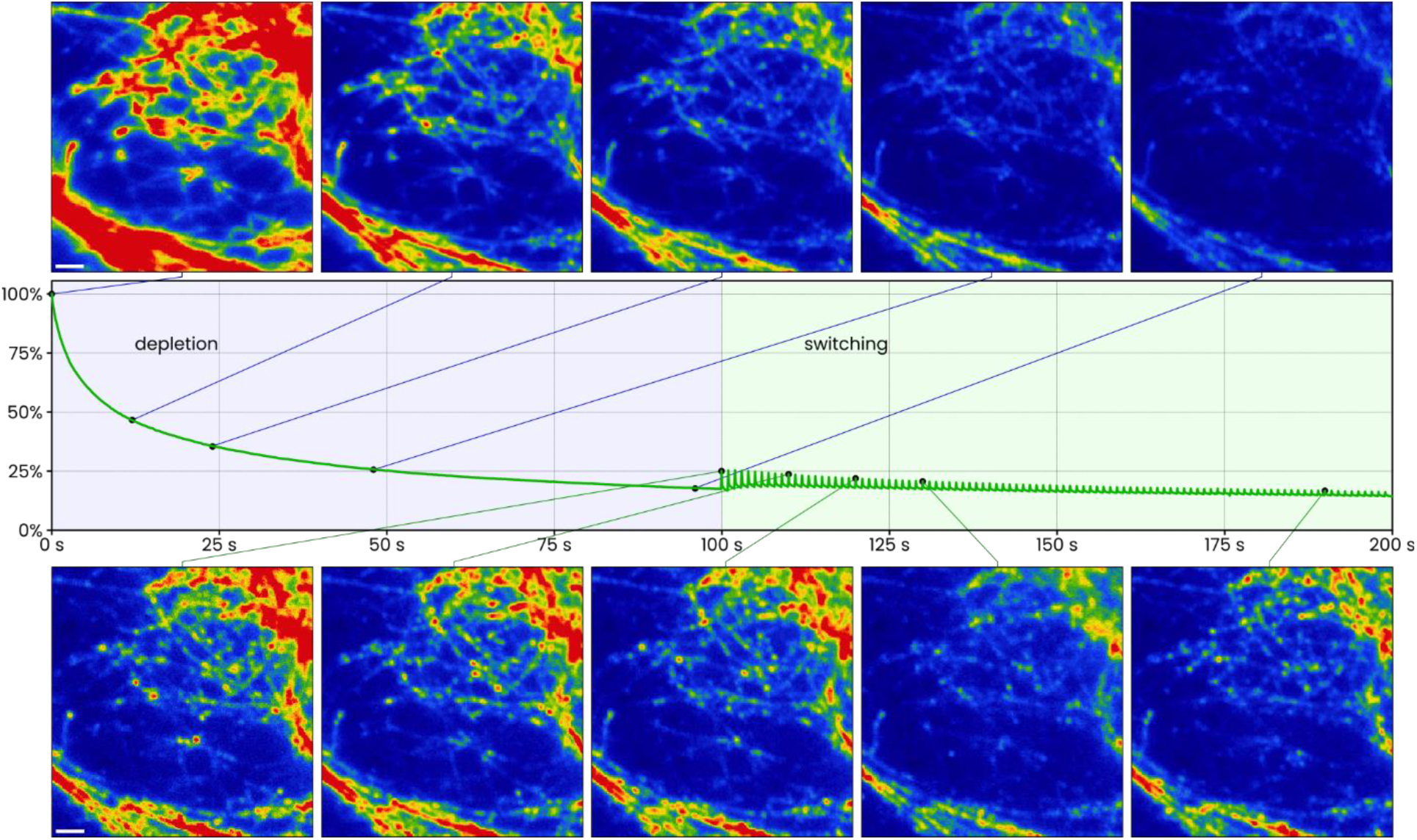
Exemplary time traces and fluorescence images for the ‘depletion’ (top) and ‘switching’ (bottom) illumination routines. For this region of the sample, an initial 100 second ‘depletion’ acquisition was used (488 nm illumination only; blue shaded region of the graph), during which the overall fluorescence was reduced to ∼ 17 % of the initial intensity. A subsequent, 100 second ‘switching’ acquisition, in which every 5 images acquired with 488 nm illumination were followed by an rsEGFP2 - activating 405 nm pulse, was then used to capture single - molecule blinking events.

**Table 1.** the ‘depletion’ acquisition routine, used to deactivate rs EGFP2 and reduce the overall fluorescence intensity level.

| Light source | Exposure time (ms) | Repeats |
| --- | --- | --- |
| 528 nm LED | 100 | 1 |
| 488 nm laser | 100 | 50 |
| *Repeated 20 times.* |  |  |

Once the overall fluorescence has been sufficiently reduced and diffraction limited single-molecule images can be observed, we employ the switching acquisition routine to acquire datasets for single-molecule localization reconstruction (Fig. 4, Table 2). In addition to 488 nm illumination for fluorescence excitation, this routine also involves an interleaved 405 nm laser illumination pulse with which rsEGFP2 is reactivated. The activation by 405 nm and deactivation by 488 nm illumination - i.e., the switching - requires some balancing: with too few 405 nm pulses the number of single-molecule switching events is low, but too many pulses can cause the overall fluorescence intensity to recover to a high value. A good starting point is to use one activation pulse per five excitation pulses, again using a total of 100 seconds of 488 nm exposure: 5 x 488 nm x 100 ms followed by 1 x 405 nm x 100 ms, repeated 200 times (∼180 W/cm^2^ and ∼100 W/cm^2^, respectively). During acquisition, the power of the 405 nm laser can be adjusted in order to tweak the photo-switching balance somewhat.

**Table 2.** the ‘switching’ acquisition routine, used to capture single-molecule blinking events. One reflection image (528 nm LED) is acquired for every 50 fluorescence images (488 nm); i.e., a reflection image is acquired, followed by ten repeats of the activation and fluorescence excitation block, and this routine is repeated 20 times.

| Light source | Exposure time (ms) | Repeats |  |
| --- | --- | --- | --- |
| 528 nm LED | 100 | 1 | A |
| 405 nm laser | 100 – 300 | 1 | B |
| 488 nm laser | 100 | 5 |  |
| Twenty rounds of: 1x A, followed by 10 x B |  |  |  |

In our experience, the data obtained using 100 seconds of acquisition is sufficient to generate single-molecule localization maps (SML maps) with enough detail to enable accurately targeted cryoET acquisition.

During every acquisition, we additionally periodically acquire reflected light images of the sample, which can later be used to guide the correlation of light and electron microscopy images. On our microscope we use a 528 nm LED light source that is partially transmitted by the rsEGFP2 fluorescence filters, with which we avoid having to change filters during an acquisition.

- Assess, based on whether single-molecule blinking events can be seen in the region of interest, if a depletion acquisition is required or not.
- If so, set up a depletion acquisition routine:
- Once completed, set up a switching acquisition routine:
- Generally, 1000 frames of the switching routine are sufficient to generate a useful SML map.

#### scNodes pre-TEM processing

We use scNodes^23^ for all of the data processing steps, except tomogram reconstruction for which we use IMOD^24^. To ensure accurate correlation of fluorescence and electron microscopy images, it is important to take the eventual registration requirements in mind from the very beginning. In this workflow we use the reflected light images to guide the registration of the light and electron microscopy datasets. These images show features that can also be recognized in EM, while they were acquired in the same reference frame as the fluorescence timelapse videos.

Data processing in scNodes is centered around a node-based processing pipeline, where every node performs a particular operation (Fig. 5a). The first step is usually to perform drift correction by grayscale registration of every image in the timelapse to a single template image. Since the final positioning of the fluorescence data is thus dependent on the positioning of that template frame, it is critical to select a template frame that is aligned with the reflected light image that will eventually guide the correlation. By selecting a fluorescence template frame that was acquired immediately after the reflected light image, this alignment is guaranteed by the rapid sequential acquisition of the two frames.

**Figure 5.**
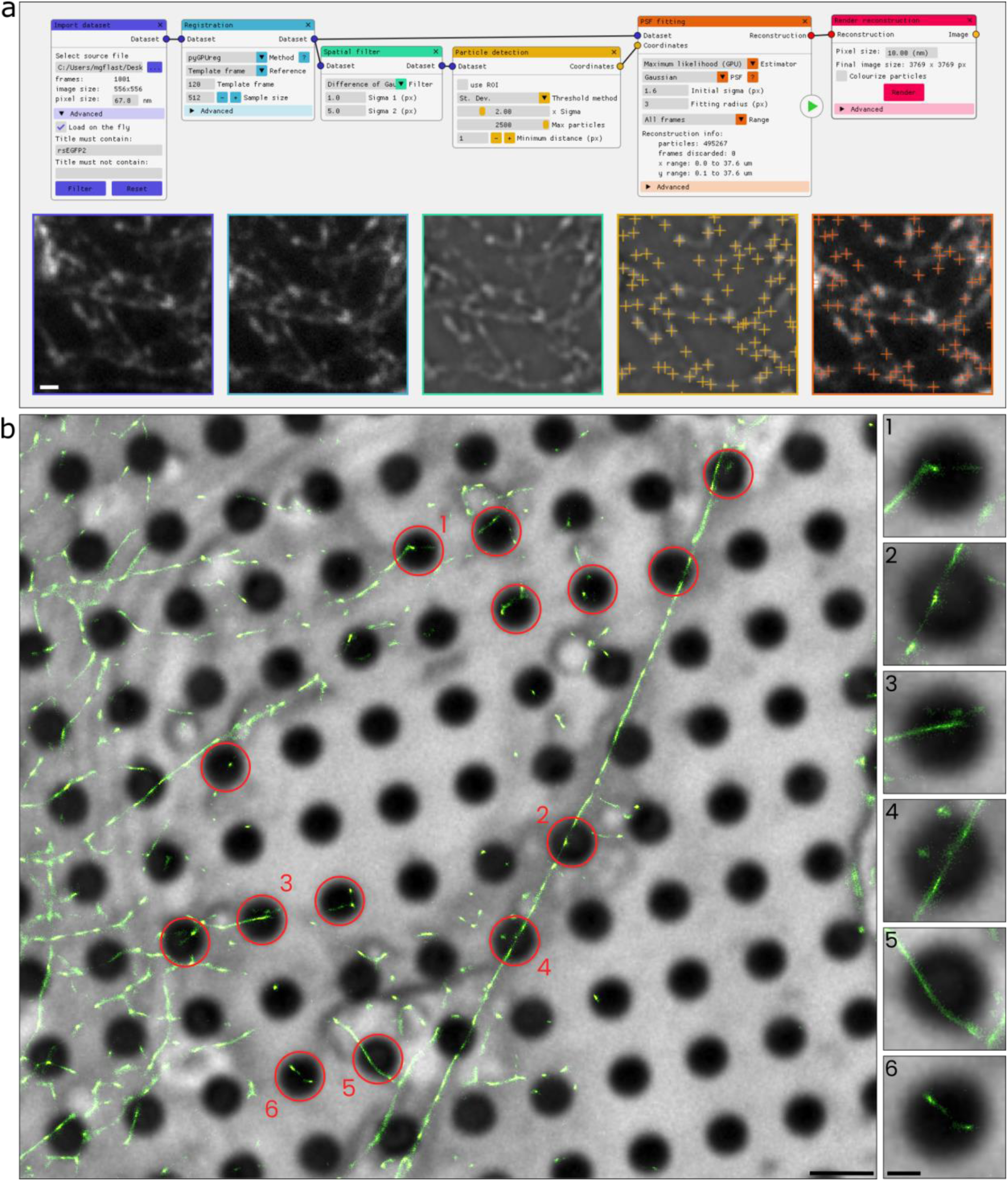
cryo SMLM results + SMLM/ reflection map for EM targeting. **a)** An overview of the scNodes node setup used to generate a single - molecule localization map based on a 100 second ‘switching’ acquisition. From left to right, the processing blocks are responsible for: loading data, drift correction by grayscale registration, background subtraction using a difference of gaussians filter, peak detection, PSF - fitting, and rendering the reconstruction. Corresponding crops of the image output of each of these nodes is displayed below in the same order (left to right). **b)** The output of the rightmost block is the SML map (green), registered to and overlayed on an unprocessed reflected light image (grayscale). A number of holes that would be of interest for tilt series acquisition are indicated in red; six of these are highlighted in the adjacent panels. In each, a filamentous (Vimentin) structure is found close to the center of the hole. Scale bars are 1 µm (a), 2 µm (b), and 500 nm (b, side panels).

For help with processing, consult the scNodes video tutorials at youtu.be/VzkF7arMnHI, or the discussion forum and manual via github.com/bionanopatterning/scNodes.

- Add a ‘Load dataset’ node and open one of the datasets that was previously acquired. If you have multiple datasets in separate folders, but want to process them as one dataset, right-click the node and select ‘Append dataset’ to concatenate multiple datasets.
- Select a reflected light image that is well-focused, right-click the image in the image browser, and select ‘Add to correlation editor’. Next, select the first fluorescence frame that was acquired after the reflected light image and add it to the correlation editor as well. This will be the template frame for drift correction.
- In the ‘Load dataset’ node, under the ‘Advanced’ header, apply appropriate filters to the dataset to remove all but the rsEGFP2 fluorescence frames.
- Add a ‘Registration’ node and connect it to the setup. In the image browser, find the index of the template frame, and enter this value in the ‘Template frame’ field of the registration node.
- Add a ‘Spatial filter’ node and set it to apply a ‘difference of gaussians’ filter (standard deviations set to 1 and 5 pixels) to the registered data. This works well for background subtraction and noise reduction, thus preparing the dataset for detecting local maxima.
- Add a ‘Particle detection’ node and input the filtered dataset. While inspecting the output of this node in the image browser, change the threshold value (or method) to an appropriate value such that single-molecule images are picked up, but noise is not.
- Add a ‘PSF fitting’ node and connect the output of the ‘Registration’ node to the dataset input, and the output of the ‘Particle detection’ node to the coordinates input. Set the ‘Range’ field to ‘Random subset’ and press the green play button to start processing a random subset of 50 frames of the input dataset.
- Add a ‘Render reconstruction’ node, connect it to the output of the ‘PSF fitting’ node, and press ‘Render’ to generate a SML map. If the resulting map does not look good (e.g. there are visible artifacts, or only few particles are fitted), adjust any of the settings in the node setup and generate a new map. It can also be helpful to focus (right-click a node, select ‘Focus’) a node: the image browser will show the output of the focused node, while you can edit the settings of any of the nodes in the setup.
- After optimizing the processing pipeline, set the range in the ‘PSF fitting’ node to ‘Entire dataset’ and start the processing. When the final SML map is complete, right-click it in the image browser and add it to the correlation editor.
- Save the node setup (File -> save setup). In the original ‘Load data’ node, you can now select any of the remaining datasets and process is using the exact same node setup.
- Open the correlation editor (main menu bar -> Editor -> Correlation Editor). The reflection image, template frame, and SML map, all of which are aligned within the same coordinate system, are visible here. Hide the template frame (select it and press the ‘V’ key, or uncheck it in the menu) and move the SML map onto the reflected light image. Set the lookup table of the localization map to green, and its blend mode to ‘Sum’ (press the ‘2’ key). The resulting overlay of single-molecule localizations on the reflected light image (Fig. 5b) can now be used to target cryoET data acquisition. Save the correlation editor scene (main menu bar -> File -> save scene)

### Targeted CryoET

#### Fluorescence-guided tilt series acquisition

The cryoET part of an SR-cryoCLEM acquisition is not remarkably different from any other cryoET experiment, except that the selection of sites to acquire tilt series at is guided by the super-resolution fluorescence images. We will thus start from the point where the sample is loaded into the microscope, the microscope is aligned, and atlases have been acquired. The nomenclature in the upcoming section is based on that used in the *Tomography 5.0* acquisition software (Thermo Fisher Scientific). On our 200 keV Talos Arctica microscope we typically use the following magnification presets: Atlas (110×), Overview (560×), Search (7300×), and Exposure (31000×, with a pixel size of 2.74 Å).

It is very useful to have access to both scNodes and the electron microscope’s data server from the same workstation, in order to rapidly load the EM data into scNodes, perform correlations, and then decide where to acquire tilt series. How this can be achieved depends on your local infastructure; we run scNodes via a remote connection to a PC that has access to our cryoEM file server.

A video tutorial for the below procedure is available at youtu.be/zAsTFK2rvI8

- Open the correlation editor in scNodes and load one (or multiple) of the previously saved scenes.
- Locate this same region of interest in the atlas image. Finding and measuring the distance to the center marker in the atlas image can be helpful if the position of the center marker and the sites of interest were also recorded during the light imaging step. Usually, it is fairly straightforward to relocate previously imaged sites, based on the morphology of the sample as seen in both the reflected light and EM atlas image. Depending on the orientation with which the samples were loaded in the microscopes, the EM images may be rotated as well as mirrored relative to the light microscopy images.
- After relocating a site of interest, first focus then set the stage to the eucentric position, and then acquire an overview image at this site. Right-click the resulting image, select ‘save without overlay’, and save it as an .mrc file on a drive that you can access with scNodes.
- Import the overview image into the scene in scNodes. Hide all images except the overview and reflected light image, move the overview image to the background, and manually register the reflected light image onto the overview image, based on the features that are visible in both image modalities: cells, and holes in the support film (Fig. 6a).
- Hide the reflected light image and overlay the SML map on the overview image (Fig. 6b). The resulting image can then be used to target tilt series acquisition. Save the correlation editor scene.
- Identify sites that are suited for acquisition, i.e., that contain the protein of interest, are sufficiently far from the grid bars to allow for imaging at high tilt angles, and are sufficiently thin. Acquire a search image at these sites, and set up these positions for batch tilt series acquisition. Optionally, save and import the search images into scNodes, and position them in the scene by aligning them with the corresponding region seen in the overview image (Fig. 6c). This preliminary view will give some idea of what structures might be visible in the final dataset. Number and mark the holes in which a tilt series will be acquired.
- Repeat the above procedure for any other grid squares, then start the tilt series acquisition. We typically use a tilt range of -51° to +51°, in a dose-symmetric acquisition scheme with 3° steps, and a total dose of 60 electrons per Å^2^, but these acquisition parameters vary based on the goal of the experiment.

**Figure 6.**
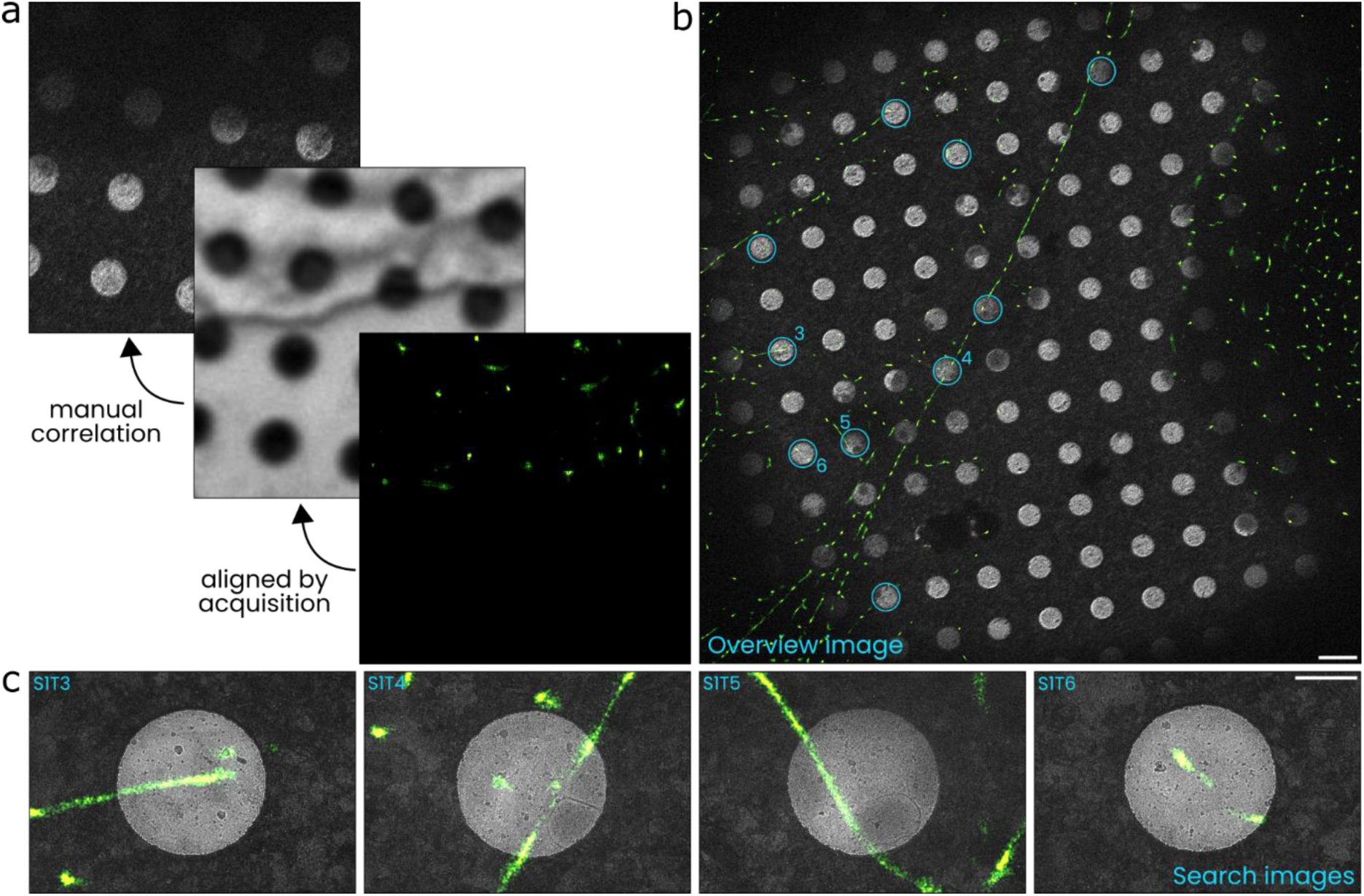
Targeted tilt series acquisition based on single - molecule localizations. **a)** To access the SML map’s information within the context of the EM, the fluorescence and electron microscopy images are first correlated by use of a reflected light image as a ‘bridge’: the reflected light image can manually be correlated with an EM overview image, while it is also registered in the same coordinate system as the SML map by virtue of rapid sequential acquisition and drift correction. **b)** The SML map overlayed on a cryo EM ‘overview’ image of the same grid square. Red circles indicate the positions of the 10 tilt series that were acquired in this region. Numbers indicate correspondence to the panels below, as well as to the ROIs in Figure 4. **c)** cryo EM ‘search’ images overlayed with the SML map, acquired during the course of setting up the batch tilt series acquisition. At this magnification, Vimentin filaments (or similar structures such as actin filaments, microtubules, etc.) can be visible but are hard to identify. Scale bars are 2 µm (b), 500 nm (c).

#### Tomogram reconstruction

To prepare the tomograms for correlation with single-molecule localization data, we process the tilt series in IMOD^24^. This procedure is described in great detail elsewhere and will not be covered here, except for one detail that is of importance for the accuracy of the final correlation. One processing step that is often applied during tomogram reconstruction is ‘tomogram positioning’; i.e., orienting the reconstruction in such a way that the bounding box is smallest. As we will use the 0° tilt image in a later correlation step, it is important not to change the orientation of the data. This would otherwise lead to the tomogram being warped relative to the 0° tilt image.

For ease of importing into scNodes and optimal segmentation results, we also do not convert the reconstruction into an integer data format, but use (32 or 16 bit) floats instead, and we re-orient the volume such that the Z axis is the first dimension (in IMOD postprocessing: rotate 90° around the X axis, or swap the X and Z dimensions).

### Correlation

#### scNodes post-TEM processing

After tomogram reconstruction, all the datasets that are required to generate the final SR-cryoCLEM dataset have been prepared. At this point, the correlation editor scenes contain the cryoEM overview image aligned to and overlayed with the SML maps, and the next step is to position the tomograms within the scene. We do so in multiple steps with increasing magnification: first, we register the search images to the overview image, then the 0° tilt images to the search image, and finally we align the tomogram with the exposure image (Fig. 7a). By thus aligning the tomograms with the overview image, which was previously correlated with the reflected light image that defined the positioning of the SML map as well, we obtain the final correlation of the tomograms with the SML map.

**Figure 7.**
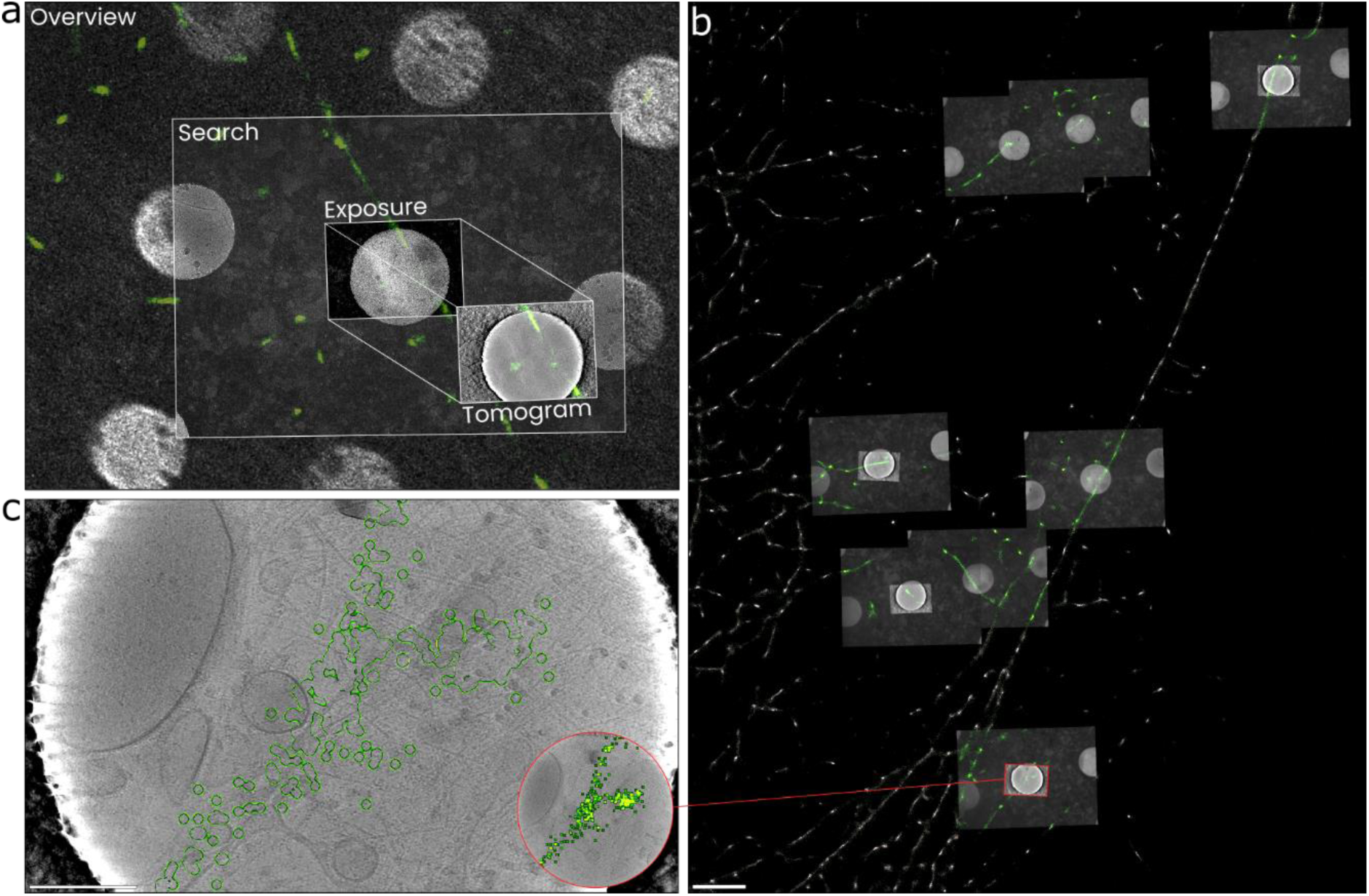
EM image alignment to obtain a final correlated cryoSMLM & cryoET dataset. **a)** EM images are aligned sequentially: search to overview, then exposure to search, then tomogram to exposure. Since the overview image was previously correlated with the SML map, the result is that the tomograms are also aligned with the SML map. In this figure, the tomogram is offset relative to its true position for illustrative purpose. **b)** A view of the original, large field-of-view SML map, overlayed with tomograms and search images. The initial manual correlation of the overview and reflected light images needs only be performed once per grid square, and can then be used for tomograms acquired in that grid square. One tomogram, highlighted in red, is shown in detail in panel c. **c)** A detailed view of one of the annotated tomograms. Fluorescent particles were located within the areas encircled in green. The inset shows the same region but with the SML map rendered in full, rather than as an outline. Scale bars are 2 µm (b) and 200 nm (c).

A video tutorial for the below procedure is also via youtu.be/zAsTFK2rvI8, and a full tutorial is included in the scNodes user manual, available at github.com/bionanopatterning/scNodes.

- In scNodes, load the correlation editor scene that contains the correlated overview image and SML map. For every tomogram that was acquired at this site, import the search image, the 0° tilt image, and the tomogram.
- Roughly position the search images over the corresponding region in the overview image, then use the ‘Register grayscale’ plugin to automatically align the search and overview image. Alternatively, align the two images manually. The various image blend modes, look-up tables, and clamping modes can be used to render the images in ways that facilitate this alignment.
- Repeat the previous procedure to align the 0° tilt images to the search images.
- Before repeating the previous procedure once more to align the tomograms to the 0° tilt images, browse the tomogram until you find a digital slice that shows features that are also visible in the 0° tilt images. Ice particles, the edges of the holes in the support film, or other high contrast features are suitable features for alignment.
- Select the SML map, right-click it, and click ‘bring to front’ to overlay it on the

tomograms (Fig. 7b). Save the scene.

- With that, the final correlated dataset is complete. The single-molecule localization annotated tomograms can now be inspected in detail in the correlation editor (Fig. 7c), and various subsequent processing steps are additionally available in scNodes; e.g., exporting high resolution images, particle picking, and segmentation.

#### Fluorescence-guided particle picking and segmentation

After this final correlation step, the correlated super-resolution fluorescence and cryo-electron tomography dataset is completed. The result is a high-resolution *in situ* map that combines an EM volume with a single-molecule localization map that pinpoints the location of the fluorescently labelled structure of interest.

Besides enabling targeted cryoET acquisition, SR-cryoCLEM thus also facilitates the identification of the structure of interest within the crowded interior of the cell, which allows fluorescence-guided particle picking (Fig. 8ab), as well as fluorescence-guided segmentation (Fig. 8cd). While outlining these applications is outside of the scope of this current chapter, we end this chapter by demonstrating these principles in the below figure, and with a discussion of the potential applications of SR-cryoCLEM.

**Figure 8.**
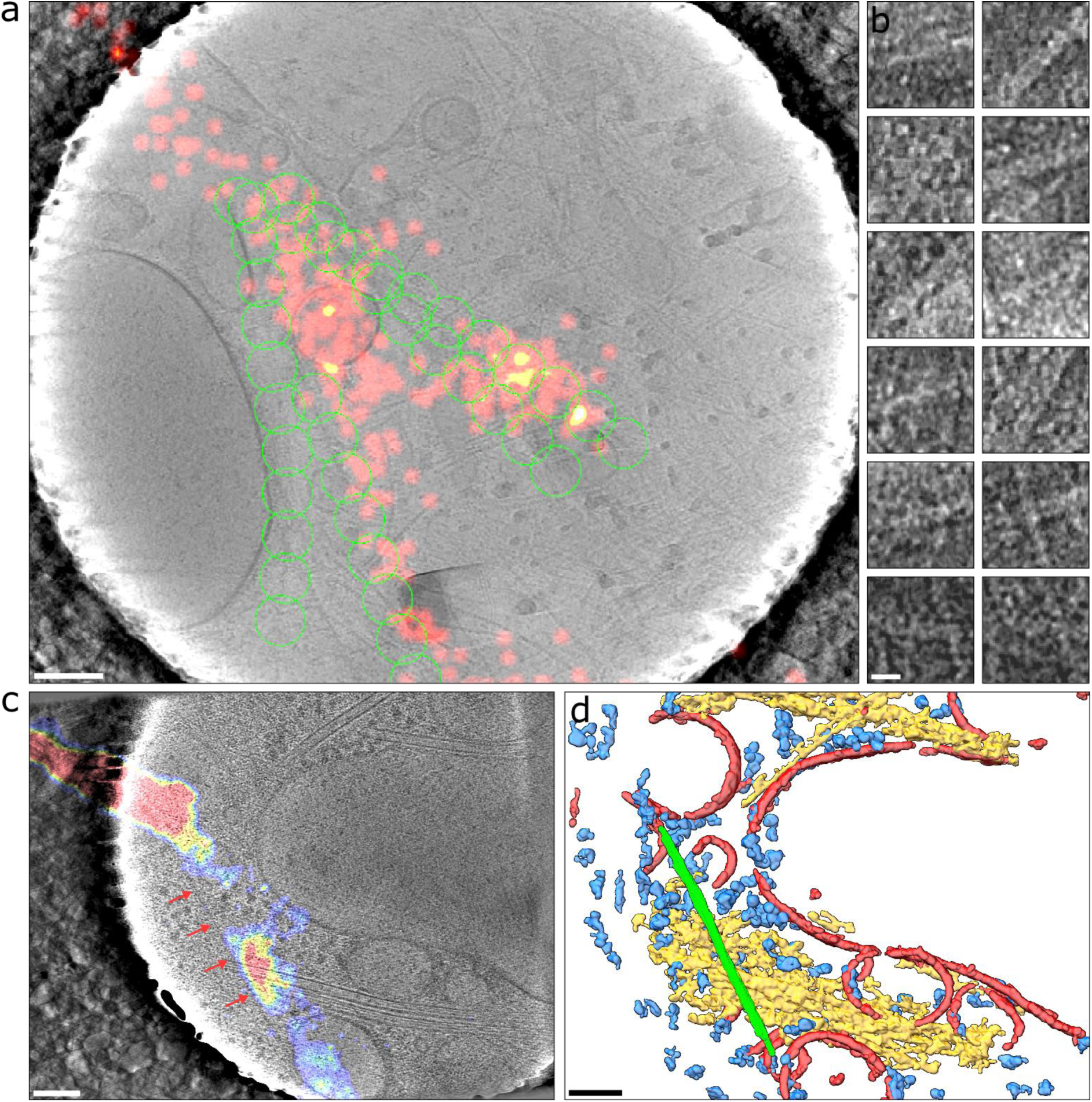
Fluorescence-guided particle picking. **a)** A tomogram overlayed with an SML map (red) and an overlay showing the location of particle ROIs, picked manually on the basis of the fluorescence localizations and the underlying tomogram morphology. **b)** A number of corresponding ROIs cropped from the tomogram. **c)** A tomogram overlayed with an SML map, renderd in heatmap colours. An approximately ∼ 11 nm thick filament can clearly be seen adjacent to the location indicated by the SML map. An ∼ 35 nm offset between the filament and the filamentous localization cloud is also apparent, and is due to a slight correlation error that occurs during the manual alignment steps. **d)** A segmentation of the tomogram in panel c. The membranes (red), microtubules (yellow), and ribosomes (blue) were segmented by convolutional neural networks, whereas the Vimentin filament segmentation was created manually. Scale bars are 100 nm (a, c, d) and 20 nm (b)

## Discussion

In this chapter we have demonstrated the full workflow of a SRcryoCLEM experiment: from sample preparation, through cryoSMLM acquisition and targeted cryoET acquisition, to post-acquisition correlation of single molecule localization maps with cryo-electron tomograms. This workflow results in high-resolution, volumetric, and fluorescence-annotated *in situ* cryoET datasets, which form the basis of downstream applications such as segmentation, to map the ultrastructure of the cell, or particle picking for protein structure determination using sub-tomogram averaging.

By outlining the steps involved in performing SRcryoCLEM as well as its potential applications, we hope to achieve two things. First, to demonstrate the SRcryoCLEM is practicable not just for those who work on method development but also to those who use cutting-edge methods for the study of biological systems. As with other developments, such as the use of FIB milling of lamellae for cryoET or the use of cryoET for bioimaging itself, popular adaptation of SRcryoCLEM and the utilization of the benefits it offers lags behind on what development has shown is feasible. Naturally, development precedes application, but in light of the high-resolution imaging opportunities that we believe SRcryoCLEM offers, we feel that it is worthwile to promote adaptation of the technique.

Secondly, we wanted to demonstrate the two opportunities that we think SRcryoCLEM offers beyond those offered by other methods of cryoCLEM that have already gained significant momentum in the field (e.g. integrated cryoFM in FIB/SEM devices). These are: 1) the accurate targeting of *in situ* tomogram acquisition, and 2) the potential to identify, based on single-molecule fluorescence localizations, biomolecules that are visible within tomograms. This enables finding and imaging rare, transient, or obfuscated cellular components, increasing the throughput of cryoET studies by significantly raising the ‘hit rate’ of data acquisition, and aiding in the interpretability of the structures seen in tomographic volumes.

There are, of course, also downsides to incorporating cryoSMLM into the cryoET workflow. As mentioned before, one is that cryoSMLM requires dedicated hardware and expertise. Recently, multiple commercial parties have released new products - microscopes, cryostages, handling tools - that facilitate cryoCLEM. In the future, we think, it would be interesting for the development of these systems to incorporate aspects of SRcryoCLEM in their design; whether via deconvolution or pixel-reassignment confocal microscopy (as in currently available cryoCLEM systems by, respectively, Leica and Zeiss), or higher-resolution approaches such as SMLM and related techniques. Widespread adoption and use of SRcryoCLEM could be greatly facilitated by the development of robust and easy to use commercial systems.

A second downside of SRcryoCLEM - and one that can similarly be resolved by the development of specialized hardware - is that of ice contamination. By introducing extra sample handling and transfer steps, significantly more contamination accumulates on a sample than when one only uses cryoEM. This is a well-known issue, yet much room for improvement remains. Again, the production of commercial cryostages that incorporate advancements recently reported by various labs^25-27^ would be a welcome development - particularly so if the sample handling interfaces of these stages are made compatible with those of widely used cryo-electron microscopes.

Development of SRcryoCLEM involves and requires advancement of more aspects than just these mentioned above: Optical systems, fluorescent probes, sample preparation methodology, software for correlation, automation, visualization, data processing - all have room for improvement, and as such, a number of advances are to be expected in the coming years: e.g., the advent of 3D and multi-colour cryoSMLM, improvement of correlation accuracy, increased resolution^28^, and higher throughput. While some of these advancements could be achieved by translating principles of ambient-condition super-resolution fluorescence microscopy into the cryogenic situation, others will be the product of the development of altogether new methodology.

In all, we believe that SRcryoCLEM is a highly promising technique that can find wide application and that is part of a field of study that is, and has historically been, very productively positioned at the interface between method development and application, and between academic exploration and commercial development. We hope that by sharing our workflow and microscope designs (in another chapter in this book), we can be of help to other groups who are interested in the application or further development of SRcryoCLEM, and we are open to any inquiries about adaptation of our methods.

## Acknowledgements

We thank Paul Verkade for their helpful feedback.

## Footnotes

1 See Last, et al. (2023) “Building a super-resolution fluorescence cryomicroscope”, available as a preprint on bioRxiv

## References

1 Guaita, M., Watters, S. C. & Loerch, S. Recent advances and current trends in cryo-electron microscopy. Current Opinion in Structural Biology 77, 102484 (2022). 10.1016/j.sbi.2022.102484

2 Jun, S. et al. Advances in Cryo-Correlative Light and Electron Microscopy: Applications for Studying Molecular and Cellular Events. The Protein Journal 38, 609–615 (2019). 10.1007/s10930-019-09856-1

3 Klein, S. et al. Post-correlation on-lamella cryo-CLEM reveals the membrane architecture of lamellar bodies. Communications Biology 4, 137 (2021). 10.1038/s42003-020-01567-z

4 Dahlberg, P. D. et al. Cryogenic single-molecule fluorescence annotations for electron tomography reveal in situ organization of key proteins in Caulobacter. Proceedings of the National Academy of Sciences 117, 13937–13944 (2020). 10.1073/pnas.2001849117

5 Tuijtel, M. W., Koster, A. J., Faas, F. G. A. & Sharp, T. H. Correlated Cryo Super-Resolution Light and Cryo-Electron Microscopy on Mammalian Cells Expressing the Fluorescent Protein rsEGFP2. Small Methods 3, 1900425 (2019). 10.1002/smtd.201900425

6 van Driel, L. F., Valentijn, J. A., Valentijn, K. M., Koning, R. I. & Koster, A. J. Tools for correlative cryo-fluorescence microscopy and cryo-electron tomography applied to whole mitochondria in human endothelial cells. European Journal of Cell Biology 88, 669–684 (2009). 10.1016/j.ejcb.2009.07.002

7 Koning, R. I., Koster, A. J. & Sharp, T. H. Advances in cryo-electron tomography for biology and medicine. Annals of Anatomy - Anatomischer Anzeiger 217, 82–96 (2018). 10.1016/j.aanat.2018.02.004

8 Berger, C. et al. Cryo-electron tomography on focused ion beam lamellae transforms structural cell biology. Nature Methods 20, 499–511 (2023). 10.1038/s41592-023-01783-5

9 Gemmer, M. et al. Visualization of translation and protein biogenesis at the ER membrane. Nature 614, 160–167 (2023). 10.1038/s41586-022-05638-5

10 Floris, D. & Kühlbrandt, W. Molecular landscape of etioplast inner membranes in higher plants. Nature Plants 7, 514–523 (2021). 10.1038/s41477-021-00896-z

11 Wu, G.-H. et al. CryoET reveals organelle phenotypes in huntington disease patient iPSC-derived and mouse primary neurons. Nature Communications 14, 692 (2023). 10.1038/s41467-023-36096-w

12 Tuijtel, M. W., Koster, A. J., Jakobs, S., Faas, F. G. A. & Sharp, T. H. Correlative cryo super-resolution light and electron microscopy on mammalian cells using fluorescent proteins. Scientific Reports 9, 1369 (2019). 10.1038/s41598-018-37728-8

13 Dahlberg, P. D. et al. Identification of PAmKate as a Red Photoactivatable Fluorescent Protein for Cryogenic Super-Resolution Imaging. Journal of the American Chemical Society 140, 12310–12313 (2018). 10.1021/jacs.8b05960

14 Sartor, A. M., Dahlberg, P. D., Perez, D. & Moerner, W. E. Characterization of mApple as a Red Fluorescent Protein for Cryogenic Single-Molecule Imaging with Turn-Off and Turn-On Active Control Mechanisms. The Journal of Physical Chemistry B 127, 2690–2700 (2023). 10.1021/acs.jpcb.2c08995

15 Kaufmann, R. et al. Super-resolution microscopy using standard fluorescent proteins in intact cells under cryo-conditions. Nano Lett 14, 4171–4175 (2014). 10.1021/nl501870p

16 Grotjohann, T. et al. rsEGFP2 enables fast RESOLFT nanoscopy of living cells. Elife 1, e00248 (2012). 10.7554/eLife.00248

17 Dahlberg, P. D., Perez, D., Hecksel, C. W., Chiu, W. & Moerner, W. E. Metallic support films reduce optical heating in cryogenic correlative light and electron tomography. Journal of Structural Biology 214, 107901 (2022).

18. Last, M. G. F., Tuijtel, M. W., Voortman, L. M. & Sharp, T. H. Selecting optimal support grids for super-resolution cryogenic correlated light and electron microscopy. Scientific Reports 13 (2023). 10.1038/s41598-023-35590-x

19 Ratz, M., Testa, I., Hell, S. W. & Jakobs, S. CRISPR/Cas9-mediated endogenous protein tagging for RESOLFT super-resolution microscopy of living human cells. Scientific Reports 5, 9592 (2015). 10.1038/srep09592

20 Hampton, C. M. et al. Correlated fluorescence microscopy and cryo-electron tomography of virus-infected or transfected mammalian cells. Nat Protoc 12, 150–167 (2017). 10.1038/nprot.2016.168

21 Carter, S. D. et al. Distinguishing signal from autofluorescence in cryogenic correlated light and electron microscopy of mammalian cells. J Struct Biol 201, 15–25 (2018). 10.1016/j.jsb.2017.10.009

22 Last, M. G. F., Voortman, L. M. & Sharp, T. H. Measuring cryo-TEM sample thickness using reflected light microscopy and machine learning. Journal of Structural Biology 215, 107965 (2023). 10.1016/j.jsb.2023.107965

23 Last, M. G. F., Voortman, L. M. & Sharp, T. H. scNodes: a correlation and processing toolkit for super-resolution fluorescence and electron microscopy. Nature Methods, 1–2 (2023). 10.1038/s41592-023-01991-z

24 Mastronarde, D. N. & Held, S. R. Automated tilt series alignment and tomographic reconstruction in IMOD. Journal of Structural Biology 197, 102–113 (2017). 10.1016/j.jsb.2016.07.011

25 Xu, X. et al. Ultra-stable super-resolution fluorescence cryo-microscopy for correlative light and electron cryo-microscopy. Science China Life Sciences 61, 1312–1319 (2018).

26 Sartori, A. et al. Correlative microscopy: Bridging the gap between fluorescence light microscopy and cryo-electron tomography. Journal of Structural Biology 160, 135–145 (2007). 10.1016/j.jsb.2007.07.011

27 Nahmani, M., Lanahan, C., DeRosier, D. & Turrigiano, G. G. High-numerical-aperture cryogenic light microscopy for increased precision of superresolution reconstructions. Proceedings of the National Academy of Sciences 114, 3832–3836 (2017). 10.1073/pnas.1618206114

28 Siegfried, W., Bo, J., Alois, R. & Vahid, S. Cryogenic localization of single molecules with angstrom precision. Proc.SPIE 8815, 88150D (2013). 10.1117/12.2025373

